# HiCDiffusion - diffusion-enhanced, transformer-based prediction of chromatin interactions from DNA sequences

**DOI:** 10.1101/2024.02.01.578389

**Authors:** Mateusz Chiliński, Dariusz Plewczynski

## Abstract

Prediction of chromatin interactions from DNA sequence has been a significant research challenge in the last couple of years. Several solutions have been proposed, most of which are based on encoder-decoder architecture, where 1D sequence is convoluted, encoded into the latent representation, and then decoded using 2D convolutions into the Hi-C pairwise chromatin spatial proximity matrix. Those methods, while obtaining high correlation scores and improved metrics, produce Hi-C matrices that are artificial - they are blurred due to the deep learning model architecture. In our study, we propose the HiCDiffusion model that addresses this problem. We first train the encoder-decoder neural network and then use it as a component of the diffusion model - where we guide the diffusion using a latent representation of the sequence, as well as the final output from the encoder-decoder. That way, we obtain the high-resolution Hi-C matrices that not only better resemble the experimental results - improving the Fréchet inception distance by an average of 12 times, with the highest improvement of 35 times - but also obtain similar classic metrics to current state-of-the-art encoder-decoder architectures used for the task.

## Introduction

With the development of the experimental methods, the spatial organisation of the chromatin became of high importance and interest to the scientific community. The spatial landscape is vital for understanding how genetic machinery works (Bouwman *et al*., 2022) - and can be used in e.g. precision medicine (Chen *et al*., 2023), for example, by improving gene expression prediction (Chiliński, Lipiński, *et al*., 2023). However, those spatial experiments are expensive and time-consuming to run. Thus, multiple efforts have been made to predict the spatial organisation of chromatin in the nucleus using various machine-learning models. The first ones relied on epigenetic signals (e.g. ChIP-Seqs, or ATAC-Seqs) or peaks (Belokopytova *et al*., 2020; Li *et al*., 2019; Whalen *et al*., 2016; Al Bkhetan and Plewczynski, 2018). However, the community wanted to use the most easily accessible genomic data, i.e. DNA sequence to identify *in silico* the spatial landscape of chromatin for specific cell lines. Multiple algorithms were proposed, providing end-to-end solutions from DNA-Seq to chromatin 3D interactions prediction. Such predictions, based only on the sequence, are of high interest because they allow the incorporation into the model genome sequence mutations that might cause changes in the spatial organisation (Chiliński *et al*., 2022), thus changing the expression or behaviour of genetic machinery significantly. To account for that, differences between the reference genome and an individual one (obtained by reads from an experiment) can be calculated. Those include Single Nucleotide Polymorphisms (SNPs), short indels (insertions or deletions below 50-100bps), and Structural Variants (SVs; insertions, deletions, duplications, copy number variations and other complex rearrangements) - and they describe how an individual differs from the reference. Applying those variants to the reference creates a personalised genome of an individual. These statistical models allow to discover the spatial landscape of chromatin for a given individual using only DNA sequence - which is relatively easy to obtain and process. Multiple bioinformatics pipelines have been proposed to identify single nucleotide mutations and structural variants from raw DNA-seq data (Chiliński and Plewczynski, 2022; Poplin *et al*., 2018; Van der Auwera and O’Connor, 2020). Those pipelines automate the whole variant discovery, taking as input in most cases either raw reads from DNA-Seq experiment (in the form of fastq files) or already-aligned data, and perform a series of calls to multiple algorithms for the variant discovery. The user is left with a list of variants, which can be used later for further analyses. We envision the entire *in silico* workflow for in-silico prediction of chromatin structure (Hi-C pairwise contact matrix) from DNA sequence, starting from DNA-Seq experimental data, running sequence variant discovery, mapping the variants to the reference genome, and running the selected 3D predictive computational model on personalised DNA sequence. Those Hi-C prediction models include convolutional networks (Cao *et al*., 2021), transfer learning-based convolutional networks (Schwessinger *et al*., 2020), convolutional encoder-decoder architectures (Zhou, 2022; Fudenberg *et al*., 2020), transformer-based methods (Chiliński, Halder, *et al*., 2023), and finally, convolutional encoder-decoder, transformer-based architectures (Tan *et al*., 2023). Those studies have achieved high metrics, including the Pearson correlation coefficient between the real and predicted heatmaps. However, while highly informative, none of those studies addressed the visual quality of the predicted heatmaps. Distinguishing between the real Hi-C matrix and one predicted by any of the presently available tools is not difficult - as they seem to be highly blurred. While those methods preserve the most critical parts of the matrices, they lack quality assessed easily by human perception.

To address this challenge, we have decided to use the recent theoretical achievements from the computer vision field. One of the many problems that computer vision is currently facing is generating images of high quality. However, in our problem, we want to strongly guide the generation in order to reflect the actual physical nature of the underlying polymer. Thus, we are primarily interested in conditional architectures that allow us to improve the quality of the Hi-C matrices, not generate random pairwise contact maps. Multiple neural network architectures have been proposed to face this problem in computer vision - Generative Adversarial Network (Goodfellow *et al*., 2020) being a prime example. However, lately, diffusion-based models (Ho *et al*., 2020) have gained tremendous popularity due to much higher performance. There have been multiple implementations, each adding some value to the architecture and allowing to obtain even higher similarity metrics. Examples include Stable Diffusion (Rombach *et al*., 2021) or DALL-E 2 (Ramesh *et al*., 2022). However, those are still used primarily for generating new images from text description - and not improving existing ones. That is why we have decided to implement the deblurring diffusion model (Kawar *et al*., 2022; Ren *et al*., 2022; Lee *et al*., 2022) to guide the network to improve the quality of the given heatmap.

Using previously established encoder-decoder architectures, we aimed to improve the predicted heatmaps’ quality using transfer learning as a core to a deblurring, diffusion-based model. We have developed the HiCDiffusion model, which provides high metrics as obtained by other encoder-decoder-based algorithms and significantly improves the predictions’ human-readable quality. Our approach makes the predicted heatmaps almost indistinguishable from the real data, which we have measured using Fréchet inception distance.

## Results

In our study, we have developed the HiCDiffusion model, which aims to improve the Fréchet inception distance (FID), thus yielding better results in terms of quality. That metric, widely used in computer vision and image generation, decreases when the quality increases. To put it into perspective when dealing with the Hi-C experiments, we have blurred an example real Hi-C map and calculated FID scores of the original and augmented data. The results are shown in **Figure 1**.

**Figure 1.**
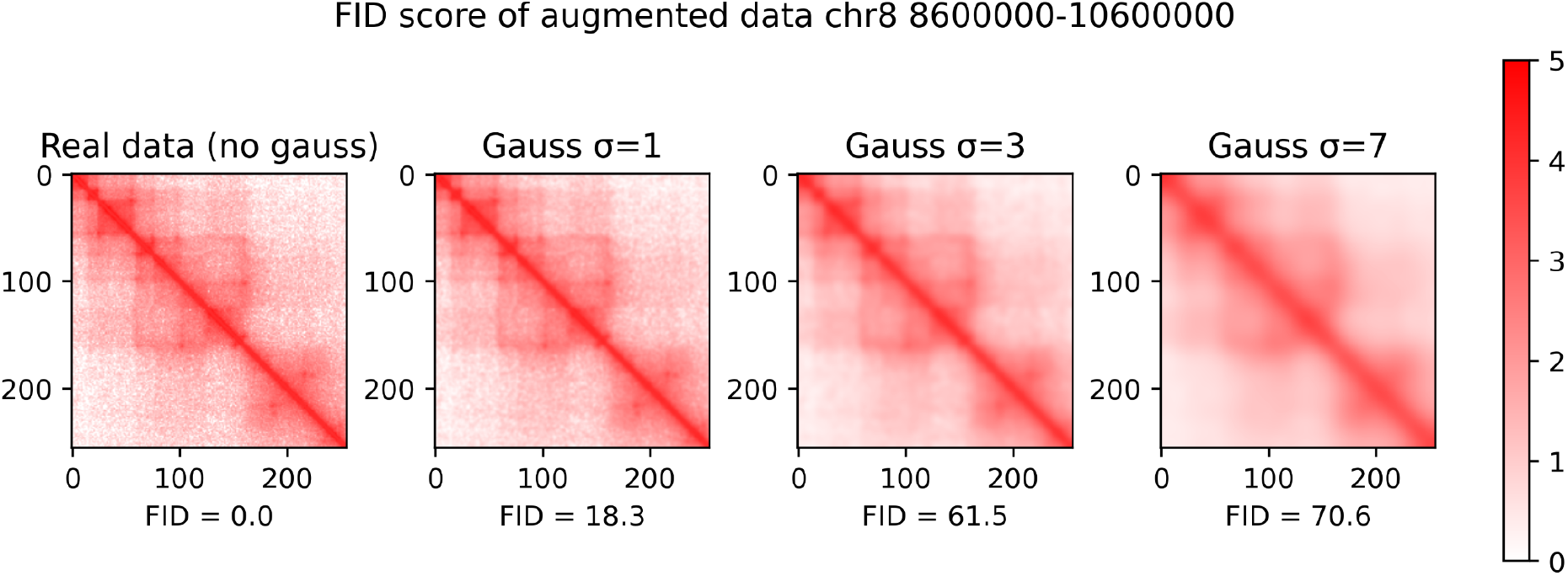
FID score of the real and augmented data - example of chr8 8 600 000-10 600 000. The first example is the real data - we compare the real data distribution with the “generated” distribution, which is also the real data. We can see that the FID score is then 0.0 - because the distributions are precisely the same. We have applied Gaussian blur to this example - in the case of blur with σ=1, the FID score increases to 18.3; in the case of the blur with σ=3, it’s 61.5, and in the last case, when Gaussian blur is performed with σ=7, we are getting FID equal to 70.6. We can clearly see that the blurring increases the FID score significantly while reducing the data quality.

In the case of FID, we deal with two distributions - one real and one generated. In the case of the situation where the real distribution is equal to the generated one, we get FID equal to 0.0 - which is the highest possible FID score. That is also the case in our example - when we take real Hi-C data and compare it to itself, we get FID equal to 0.0. The following steps are blurring the Hi-C matrix and comparing the real distribution (composed of the real data), with the blurred matrix. We have shown 3 examples with σ = (1, 3, 7). In the first case, where the blur is not that intense, as we are using σ=1, the FID score equals 18.3. In the case of the higher blur, with σ=3, we are getting a much more augmented matrix - and the FID score rises to 61.5. In the last example, we have used Gaussian blur with σ=7, and the FID score is 70.6. We can clearly see that, indeed, with the higher blurring of the matrix, we are getting a higher FID score. That is why, in our study, the goal was to decrease the FID score, that is, to deblur the Hi-C matrix generated by convolutional encoder-decoder architectures.

The Hi-C Diffusion model that we propose is composed of multiple components (see **Figure 2**). The first part, encoder-decoder architecture, is similar to the current state-of-the-art tools (Tan *et al*., 2023; Fudenberg *et al*., 2020). The input to the network is a genomic sequence - one-hot encoded, and the final output is the Hi-C matrix. We first use an encoder composed of residual 1D convolutions that transfer 1D sequence into latent space; furthermore, we use a transformer encoder to allow the model to learn long-range context. Such latent representation is then cast into the 2D matrix, and the decoder, composed of 2D convolutions with exponentially growing dilation, produces the final matrix. The last part of the encoder-decoder architecture is a convolution that transfers the latent space’s final representation into a classic Hi-C matrix of size 256×256 with one channel. The second network, which uses transfer learning (by taking pre-trained encoder-decoder architecture), is the diffusion model. Based on the previous findings about diffusion networks, we have decided that the input to the network will be residual between the real Hi-C map and the final prediction of the encoder-decoder network. Then, we apply Gaussian noise to the residual Hi-C map, and the denoising U-Net is trained to predict the noise, making it easy to obtain the actual residual Hi-C (by subtracting predicted noise from the noised residual Hi-C). The network takes as input the noised residual (or, in case of inference - the random input) and the latent representation of the Hi-C heatmap predicted by encoder-decoder architecture. Such an approach is necessary to guide the diffusion, thus creating a conditional diffusion model. For more information on the technical details of the architecture, see **Methods** and **Figure 2**.

**Figure 2.**
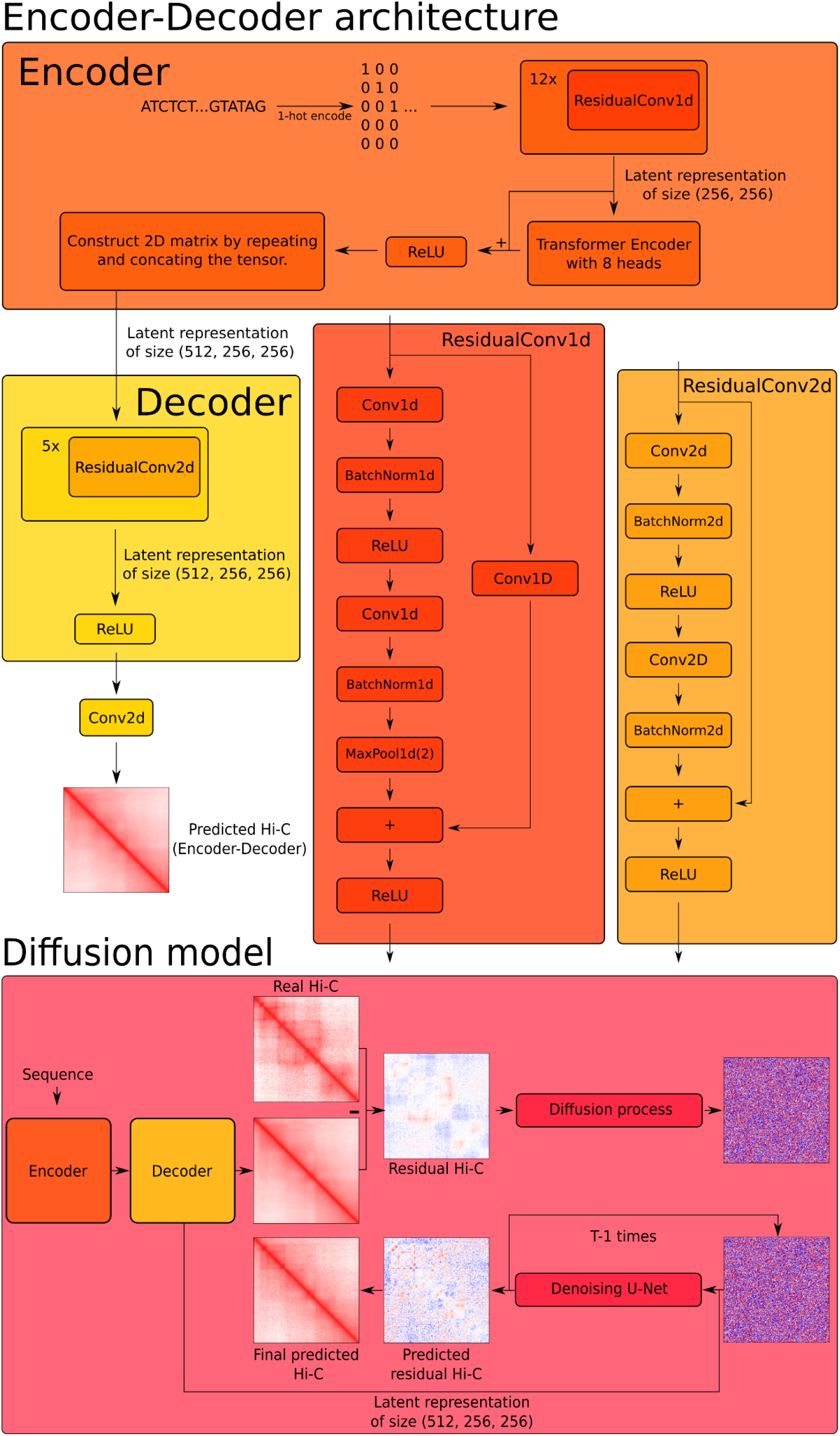
The architecture of HiC Diffusion. The example used to visualise the prediction & real data is chr8, position 8 600 000-10 600 000.

In our study, we have decided to use the context of 2,097,152 nucleotides of sequence and predict the same Hi-C region. To validate the model thoroughly, we have performed 22-fold cross-validation - creating 22 models, each with one chromosome excluded for testing purposes (see **Methods**). We have calculated the Pearson correlation coefficients for each of the examples in the testing set (for each model) and the FID score. To compare ourselves with the current state-of-the-art tool, C.Origami, we have trained the model ourselves, according to the authors’ recommendations; we have used two approaches - in one, we have excluded the epigenetic signal that they used - to keep the results consistent with our findings (see **Methods**), and second one, with epigenetic signal that they presented as the final model. We calculated and compared the Pearson correlation coefficient for those models to our model. The results are consistent and very similar - as in our work, we were aiming to obtain similar metrics in terms of correlation, in our case, even more challenging, i.e. without epigenomic profiles used as an additional input apart from the DNA sequence. Then, we calculated the FID scores for all the datasets obtained using HiCDiffusion and C.Origami. We have obtained an average improvement of FID score by 12 times in case of comparison between sequence-only models - with the highest improvement in chr7 (by 35 times). In case of comparison of our sequence-only model to C.Origami enhanced with epigenetics, the average improvement of FID score was by 11 times, and the highest improvement was also obtained in chr7 (by 32 times). The visualisation of those results can be seen in **Figure 3**, and detailed per-chromosome statistics can be found in **Supplementary Figures 1** and **2**.

**Figure 3.**
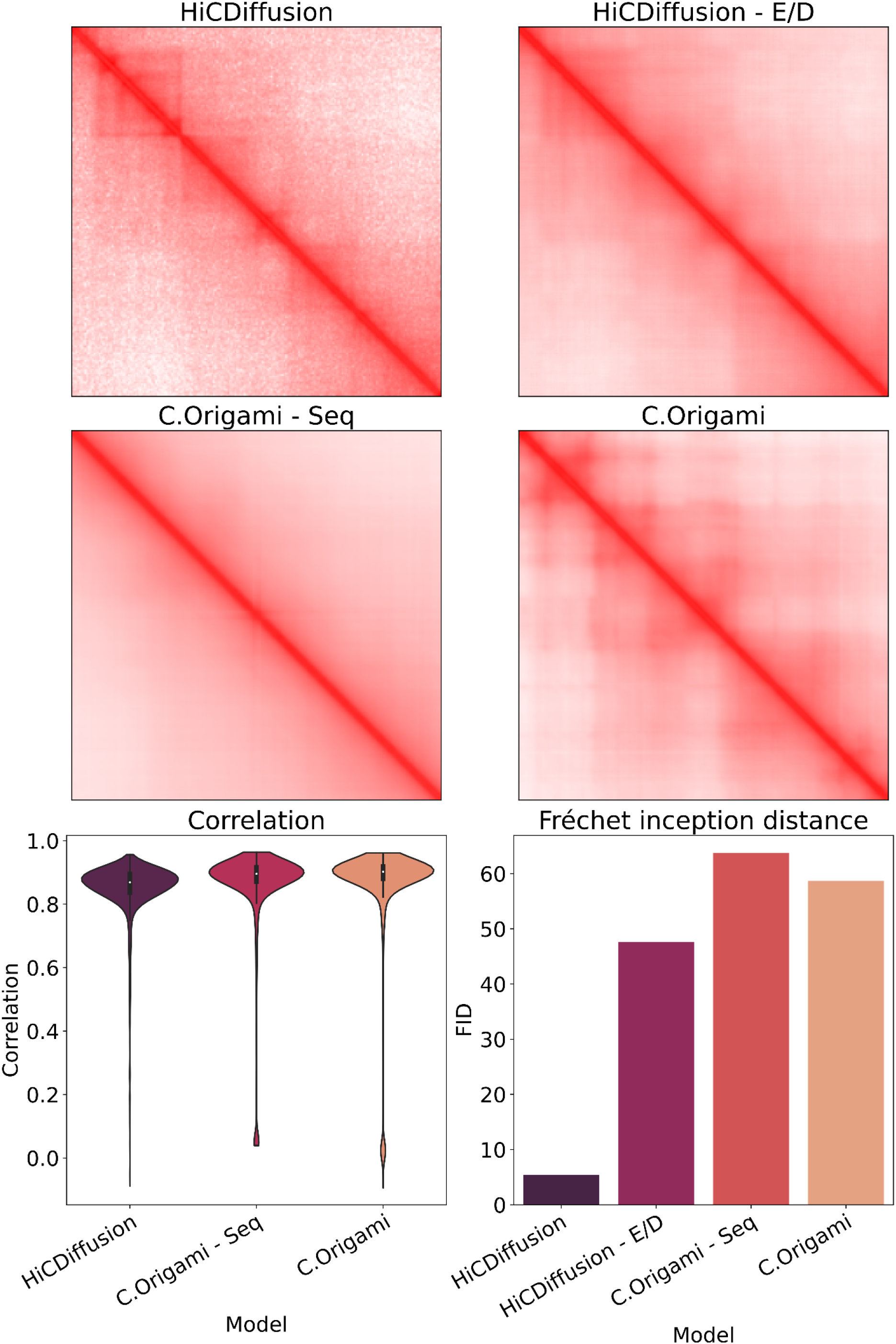
In the upper part of the figure, an example output of the 3 models is presented - full HiCDiffusion, HiCDiffusion - only encoder and decoder, and C.Origami (version with epigenetics and sequence, and version with only sequence). All example heatmaps are chr8, position 8 600 000-10 600 000. The lower part of the chart presents the Pearson correlation coefficient, based on data from all chromosomes, and the second part presents the average FID obtained by the models (the average is taken from the per-chromosome metric).

## Methods

### Data processing

The first step of processing the data is creating the genomic windows analysed in the study. The sliding window that is used for the processing of the chromosomes is set to 500kbp. We load the reference genome (GRCh38) and use pyranges (Stovner and Sætrom, 2020) to exclude telomeres and centromeres from the analysis. Then, the sequence is onehot encoded. The Hi-C matrices used in this research are taken from C.Origami (Tan *et al*., 2023) paper - we used GM12878 (Rao *et al*., 2014) cell line to allow us to compare our findings with that current state-of-the-art tool. We take precisely 2,097,152 base pairs of sequence and predict the Hi-C matrix of the same region (resized to 256×256 region) - the resolutions were chosen to easily and straightforwardly use convolutions in the network architecture.

### Training, testing, and validation data

To show the predictive power of the method, we divided the dataset into training (used for training), validation (used for choosing the best models - both encoder/decoder and diffusion), and testing (separate, used only for final testing) datasets. This division ensures that the deep learning model is generalising well and that we are unbiased toward examples occurring in the training data. To test the model entirely, we decided to create 22 models - in each, the training, validation, and testing datasets are composed of different chromosomes. For each case, we take chromosome i as the testing chromosome, chromosome i+1 as the validation, and the remaining chromosomes compose the training dataset. In the case of testing the last chromosome, chr22, the validation chromosome is chr21. We excluded sex chromosomes from the analysis.

### Architecture of the model

The architecture of the model uses the concept of transfer learning. Firstly, we use encoder-decoder architecture very similar to the ones previously published - e.g. in C.Origami (Tan *et al*., 2023) or Akita (Fudenberg *et al*., 2020). The encoder first converts the 1D genomic sequence into a sequence of 256, with 256 channels. That is done using 13 residual blocks, out of which each is composed of convolution (converting input channels into output channels, with the kernel of size 3 and padding of size 1), batch normalisation, ReLU function, another convolution (this time preserving the number of channels, with kernel of size 3, and padding of size 1), batch normalisation, and maxpooling. Additionally, the initial data provided to the residual block is downscaled using convolution (converting input channels directly into output channels, with the kernel of size 3, and padding of size 1). That downscaled data is added to the result from the previously explained sequence of transformations. The final step of the residual block is applying the ReLU function to the output. The input channels of the residual blocks used in the encoder are: (5, 32, 32, 32, 64, 64, 64, 128, 128, 256, 256, 256, 256), and the output channels are: (32, 32, 32, 64, 64, 64, 128, 128, 256, 256, 256, 256, 256). The encoder has one additional layer - the transformer encoder - that helps to learn the appropriate context of the latent representation produced by convolutions. The transformer encoder used in our study uses 8 heads. However, we also use residual connections around the transformer to preserve original data and help with the vanishing gradient problem. Finally, we apply ReLU to the output and expand the result to the 2D matrix with 512 dimensions - by repeating the (256, 256) output vertically and horizontally. That procedure leaves us with two outputs of (256, 256, 256) - one for repeating vertically and one for horizontally. We further concatenate it to the final output of the encoder of (512, 256, 256).

The decoder architecture is composed of 2D residual convolutions - each of which is composed of 2D convolution (with 512 input and output channels, kernel size of 3, and padding of 1), batch normalisation, ReLU function, and again exactly the same 2D convolution, and batch normalisation. The output of the 2D residual block is created by applying the ReLU function to the sum of the output of the previously explained sequence and the input to the residual block - this time, unlikely in the Encoder, without any downscaling. It is crucial in the decoder to use various options for the dilation parameter - it has been done in the previously mentioned papers. It is also done in our work to propagate information through the whole heatmap. For each next layer, the dilation parameter is set to (2, 4, 8, 16, 32) and represents exponential growth. Finally, after the decoder is done, we apply an additional convolutional layer (kernel size of 3 and padding of 1) that changes dimensions from 512 to 1, leaving us with the heatmap of size (256, 256).

The previously explained encoder-decoder architecture is then used for further processing by using transfer learning - we pre-train the encoder-decoder and use it for the final architecture based on diffusion models. The final architecture first computes the latent representation (output of the encoder & decoder - with 512 channel), and the final heatmap from the encoder-decoder architecture. We compute the residual heatmap by taking the difference between the real heatmap and the one produced by the encoder-decoder architecture. The network’s target is predicting the residual heatmap, which shows us how to improve the prediction of the encoder-decoder architecture. The diffusion models work in two steps - the first is the model’s training, and the second is inference. The training comes first so that it will be explained first as well. We take the residual heatmap and apply Gaussian noise multiple times - up to T. Then, we take such a noised image and use a denoising U-Net network. We try to predict the noise added to the image. The implementation of the diffusion model is pytorch reimplementation (Wang, 2020) of the original paper (Ho *et al*., 2020). We have made a few changes to make it work with the Hi-C data. Firstly, the U-Net is conditioned by taking latent encoder-decoder output (512, 256, 256). It is then converted in the initial phase of using U-Net to (32, 256, 256) using convolution with a kernel equal to 7, and padding of 3. The original image provided to U-Net (noised image) is also processed using convolution to sizes of (32, 256, 256). The condition and noised image upscaled to 32 channels are then concatenated and fed into the network to represent the conditional diffusion process. After training the diffusion model, we can predict the residual Hi-C matrix using inference mode - that is, in the form of guided diffusion using encoder-decoder output from random noise, which allows us to obtain more realistically looking Hi-C matrices.

### Testing the models

All the models were tested using independent chromosomes not used in the training or validation procedure. We have calculated the Pearson correlation coefficient for each example and Fréchet inception distance to compare the quality of the images. We have run a version of C.Origami that includes only sequence (without CTCF and ATAC-Seq signals), and compared Pearson correlation coefficient scores and FID scores.

### Fréchet inception distance (FID)

Fréchet inception distance is a metric used most often in computer vision to measure the quality of the images. It is formally defined (Fréchet and Sur, 1957) as: 

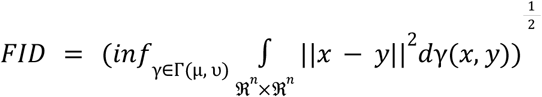

Where Γ(μ, υ)is the set of all measures on ℜ^*n*^×ℜ^*n*^ with μ as first marginal factor, and υ as the second. In our work, we are using torchmetrics (Detlefsen *et al*., 2022), which solves the aforementioned equation for multidimensional Gaussian distributions as:

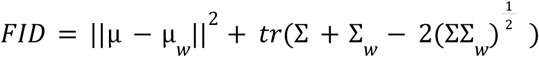

In which μ is the mean, and Σ is the variance of a multivariate normal distribution estimated from Inception v3 (Szegedy *et al*., 2015) features calculated on real data (true Hi-C matrices) and μ _*w*_ and Σ_*w*_ are corresponding values for the multivariate normal distribution estimated on the generated data (predicted Hi-C matrices). Since this metric is taking images, we need to normalise our output to the [0, 1] range and add 2 additional channels - which is done by repeating the data. The output of the procedure is called the FID score, and the lower it is, the higher quality the image is - as it is comparing the distributions of the generated images to the real ones (or, in our case, Hi-C matrices).

### Comparisons to other tools

To compare the results to the C.Origami, the current state-of-the-art tool for predicting 3D interactions, we have downloaded the original software with the dataset. However, we have used two comparison models since our study focuses on pure sequence-to-interaction relationships. In the first one, we modified the tool to accept DNA sequence and train only on it. The second one was the full C.Origami architecture - the one that includes CTCF and ATAC-Seq epigenetic signals. Then, we trained both C.Origami models, with validation chromosome chr10, and testing chromosome chr15. This approach allowed us to get accurate metrics for testing and validation chromosomes and the supremum of the metrics in the case of training chromosomes. Furthermore, this model was used to predict precisely the same windows as in our models. We have calculated the Pearson correlation coefficient for all the predictions and FID scores for all chromosomes.

## Discussion

In our study, we have created a novel deep learning architecture, HiCDiffusion, that is based on the latest advances in the field of computer vision and Artificial Intelligence. The model, composed of an encoder, transformer encoder, decoder, and a diffusion network, has surpassed the quality metric - FID score on average by 11 times (sequence-only comparison; by 10 times in case of C.Origami with epigenetics), while in the best-case chromosome, improvement was 35 times (sequence-only comparison as well; 32 in case of epigenetics-enhanced C.Origami version), over the current state-of-the-art tool, C.Origami. The level of the artificiality of the *in silico* HiC images (true quality) can also be seen with the human eye. In the case of the current models, like the aforementioned C.Origami or Akita, the model output is blurred. It can be easily distinguished from the real data. In the case of our model, we have obtained quality that can be easily confused with the experimental data while obtaining all the metrics that have made previously proposed models great. Our method is the next step towards obtaining a reliable and functional universal predictor of the spatial organisation of the chromatin within the nucleus, using purely DNA sequence - which would make connecting population studies with 3D genomics much easier and, foremost - cheaper.

## Data availability

The GM12878 Hi-C data used for comparison with C.Origami and testing of the method is available under GEO accession number GSE63525. The preprocessing steps are described in the C.Origami paper.

## Code availability

The code of HiCDiffusion is available at https://github.com/SFGLab/HiCDiffusion

