## Supplementary Figures 1 and 2 for "HiCDiffusion - diffusion-enhanced, transformer-based prediction of chromatin interactions from DNA sequences"

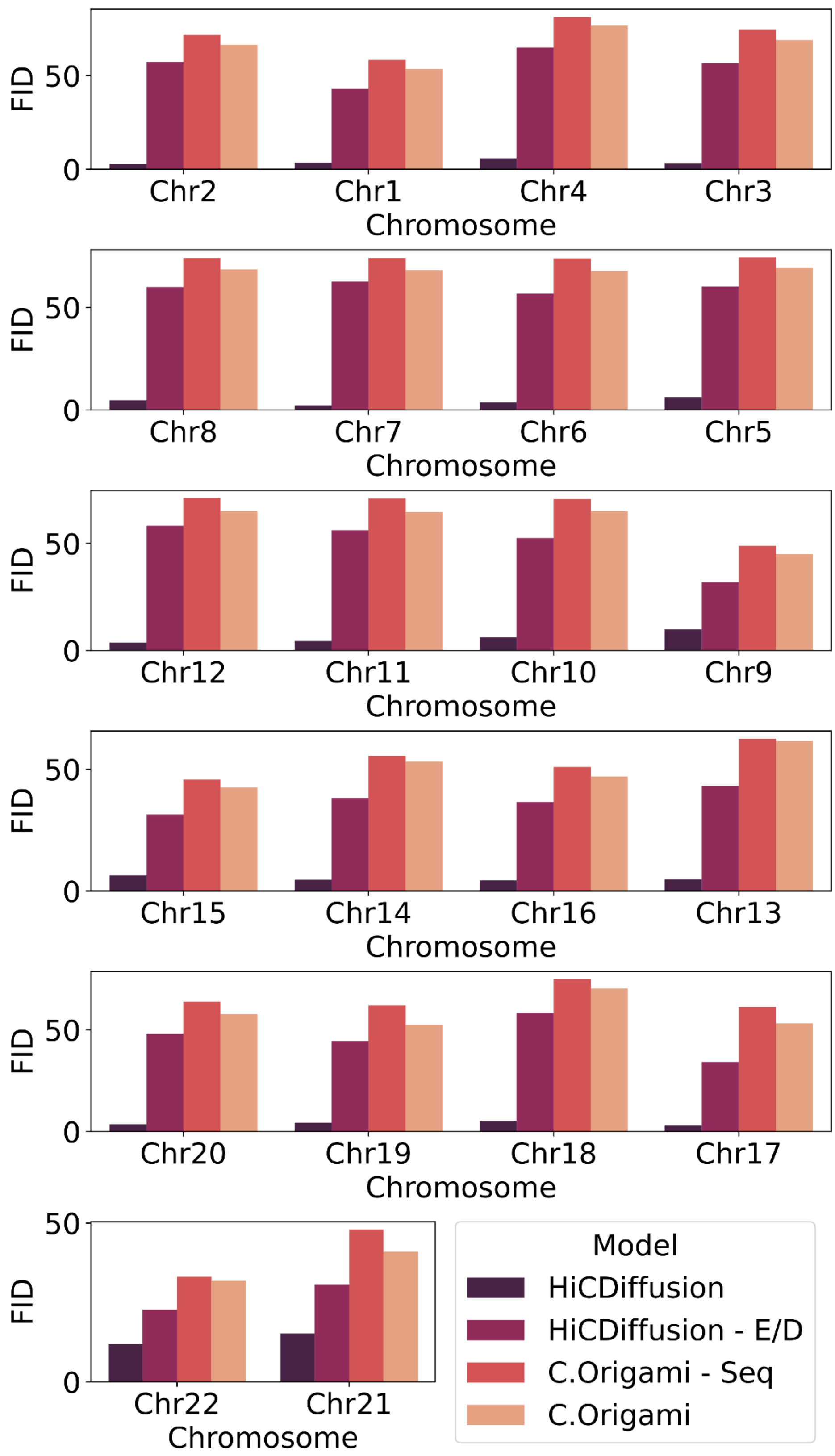


**Supplementary Figure 1.** FID score calculated for the chromosomes. In the case of HiCDiffusion, each of the charts is tied to a different model that excluded the given chromosome in validation and training data. In the case of C.Origami, one standard model (one with and one without epigenetics) was used.

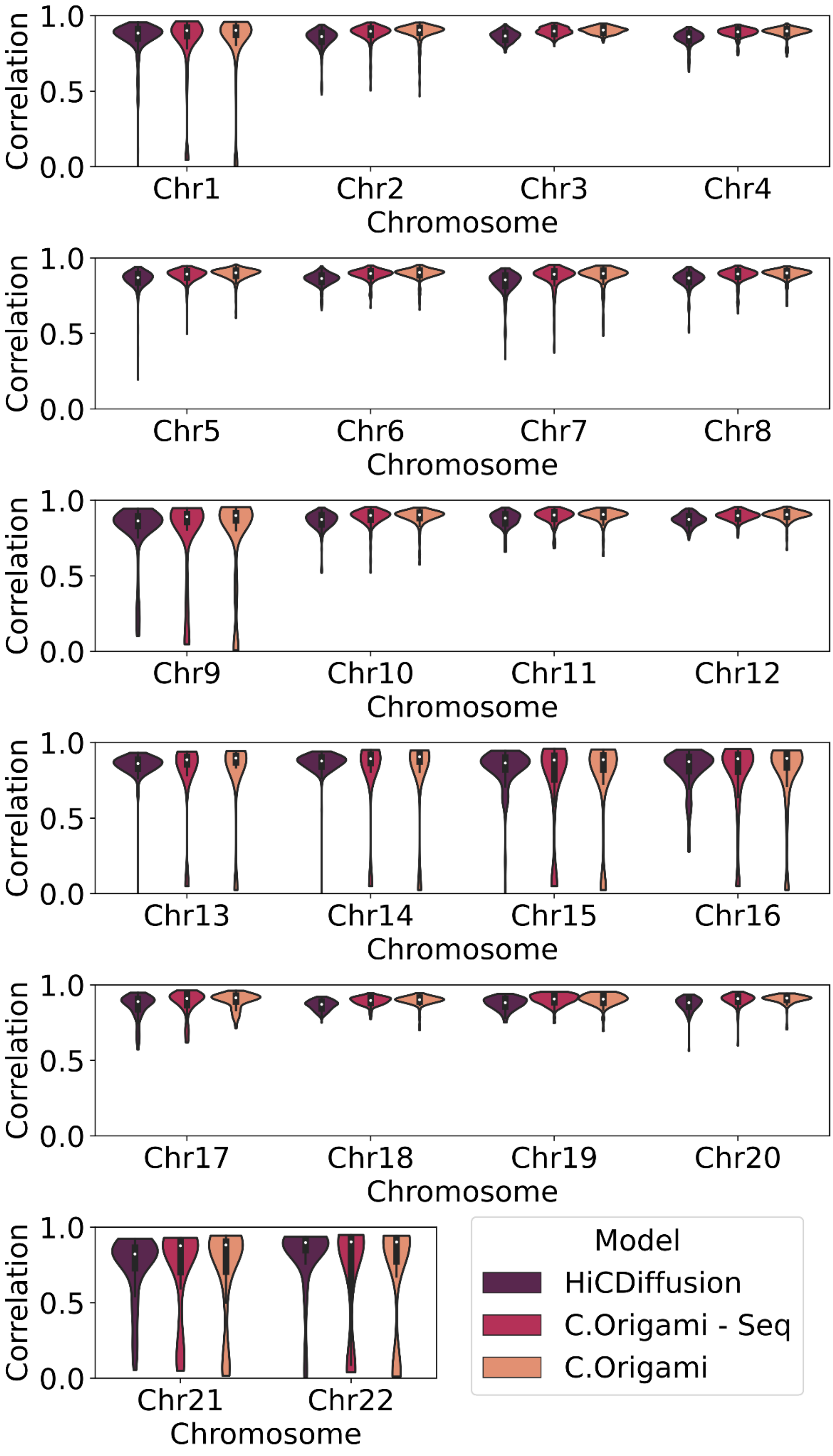


**Supplementary Figure 2.** Pearson correlation coefficient calculated for the chromosomes. In the case of HiCDiffusion, each of the charts is tied to a different model that excluded the given chromosome in validation and training data. In the case of C.Origami, one standard model (one with and one without epigenetics) was used.
